## Supplementary material for "Target protein identification in live cells and organisms with a non-diffusive proximity tagging system": POST-IT_eLife_supporting info_clean revised_v1.pdf

**This PDF file includes:**

Supplementary Methods
Figures—figure supplement
Figure 1—figure supplement 1.
Figure 1—figure supplement 2.
Figure 1—figure supplement 3.
Figure 2—figure supplement 1.
Figure 2—figure supplement 2.
Figure 3—figure supplement 1.
Figure 3—figure supplement 2.
Figure 3—figure supplement 3.
Figure 4—figure supplement 1.
Figure 4—figure supplement 2.
Figure 6—figure supplement 1.
Figure 6—figure supplement 2.
Figure 6—figure supplement 3.
Figure 6—figure supplement 4.
Data S1 to S2
References

**Other Supplementary Materials for this manuscript include the following:**

Data S1 to S2

### Supplementary Methods

#### General materials and methods for chemical synthesis

All reagents and solvents were obtained from commercial sources and used without further purification. The chemical reactions were monitored by thin-layer chromatography (TLC). $^{1}H$ and $^{13}C$ NMR spectra were recorded on AV-400 spectrometer (Bruker instrument, USA). HRMS measurements were performed with an Agilent QTOF 6520 mass spectrometer (Agilent Technologies, USA) with electrospray ionization (ESI) as the ion source. The purity of all target compounds was analyzed by HPLC (254 nm wavelength in an Agilent LC-1220 instrument) via a C18 column (5 μm, 4.6 mm × 150 mm).

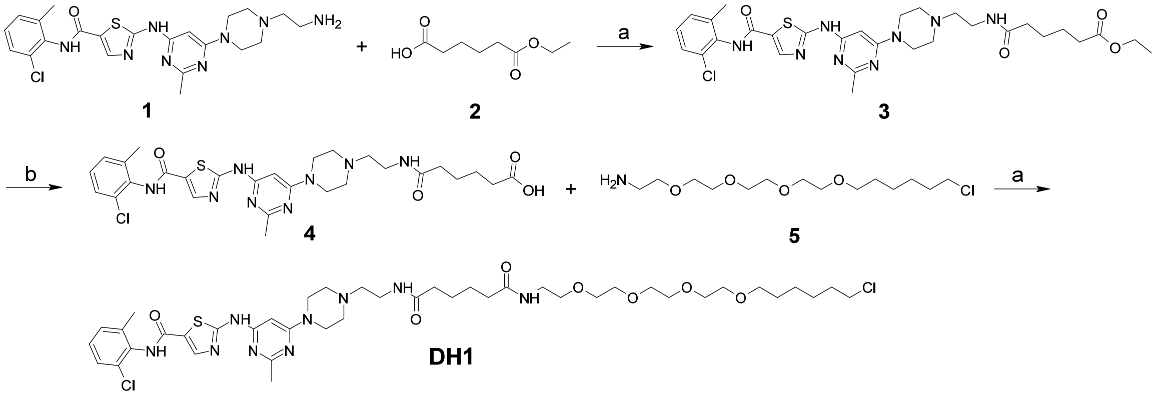


**Synthetic Scheme 1.** Synthesis of dasatinib-Halo derivative 1 (**DH1**). Reagents and conditions: (a) HATU, DIPEA, DMF, RT, 2 h; (b) NaOH, THF, $H_2O$, RT, 2 h. RT, room temperature.

*Ethyl 6-((2-(4-(6-((5-((2-chloro-6-methylphenyl)carbamoyl)thiazol-2-yl)amino)-2-methylpyrimidin-4-yl)piperazin-1-yl)ethyl)amino)-6-oxohexanoate* (**3**). A mixture of compound **1** (350 mg, 0.72 mmol), compound **2** (150 mg, 0.86 mmol), HATU (328 mg, 0.86 mmol), and 1 mL of DIPEA in DMF (10 mL) was stirred at room temperature for 2 h. The mixture was diluted with $H_2O$ (20 mL) and extracted with EtOAc (20 mL × 3). The organic layer was dried and concentrated. The residue was purified by column chromatography (DCM:MeOH = 20:1 to 15:1) to yield a white solid 210 mg. Yield: 45%. $^{1}H$ NMR (400 MHz, DMSO-$d_6$) δ 11.59 (s, 1H), 10.08 (s, 1H), 8.33 (s, 1H), 7.91 (s, 1H), 7.39 (d, $J$ = 7.6 Hz, 1H), 7.27-7.21 (m, 1H), 6.13 (s, 1H), 4.03 (q, $J$ = 7.1 Hz, 2H), 3.55-3.44 (m, 4H), 3.27-3.15 (m, 2H), 2.49-2.33 (m, 9H), 2.31-2.22 (m, 5H), 2.11-2.03 (m, 2H), 1.53-1.44 (m, 4H), 1.16 (t, $J$ = 7.1 Hz, 3H).

*N1-(2-(4-(6-((5-((2-Chloro-6-methylphenyl)carbamoyl)thiazol-2-yl)amino)-2-methylpyrimidin-4-yl)piperazin-1-yl)ethyl)-N6-(18-chloro-3,6,9,12-tetraoxaoctadecyl)adipamide* (**DH1**). To a mixture of compound **3** (141 mg, 0.22 mmol) in THF : $H_2O$ (2:1, 5 mL), NaOH was added (20 mg). The mixture was stirred at room temperature for 2 h, then concentrated under vacuum to afford compound **4**, which was used in the next step without further purification. A mixture of compound **4** (0.22 mmol), compound **7** (0.53 mmol), HATU (100 mg, 0.27 mmol), and 0.2 mL DIPEA in DCM:DMF (2:1, 9 mL) was stirred at room temperature for 2 h. The mixture was diluted with $H_2O$ (20 mL) and extracted with DCM (20 mL × 3). The organic layers were dried and concentrated. The residue was purified by column chromatography (DCM: MeOH = 10:1 to 5:1) to yield a white solid 23 mg. Yield: 11%. $^{1}H$ NMR (400 MHz, Methanol-$d_4$) δ 8.58 (d, $J$ = 4.4 Hz, 1H), 8.27 (d, $J$ = 8.0 Hz, 1H), 8.18 (s, 1H), 7.39 (dd, $J$ = 13.8, 5.5 Hz, 2H), 7.26 (q, $J$ =

8.0 Hz, 2H), 6.06 (s, 1H), 3.77-3.71 (m, 4H), 3.70-3.44 (m, 18H), 3.37 (t, $J$ = 5.3 Hz, 2H), 2.80 (s, 4H), 2.84-2.76 (m, 2H), 2.76-2.70 (s, 3H), 2.41-2.18 (m, 9H), 1.81-1.72 (m, 2H), 1.69-1.56 (m, 6H), 1.52-1.39 (m, 4H), 1.24-1.12 (m, 2H). HRMS (ESI) calcd for $[C_{42}H_{63}Cl_2N_9O_7SNa]^+$ $[M + Na]^+$, 930.3846; found 930.3843.

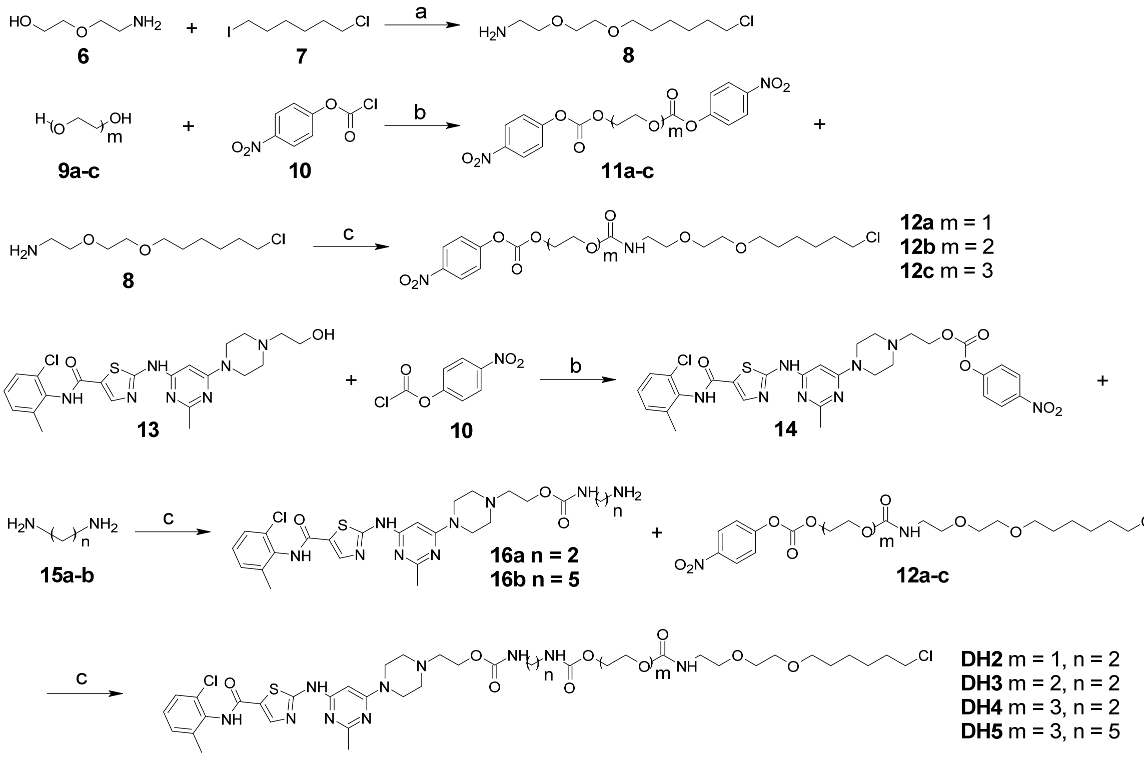

**Synthetic Scheme 2.** Synthesis of dasatinib-Halo derivatives 2-5 (**DH2-5**). Reagents and conditions: (a) NaH, DMF, 0 °C, 2 h; (b) $Et_3N$, DCM, RT, 12 h; (c) DMF, RT, 2 h. RT, room temperature.

*2-(2-((6-Chlorohexyl)oxy)ethoxy)ethan-1-amine* (**8**). A mixture of compound **6** (2 g, 19.02 mmol) in DMF (10 mL) was added to NaH (1.14 g, 28.53 mmol) and stirred at 0 °C for 30 min. Then, compound **7** (5.6 g, 22.82 mmol) in 10 mL DMF was added slowly. The mixture was stirred at 0 °C until the reaction was complete, as monitored by TLC. The reaction was quenched by adding 40 mL of saturated $NH_4Cl$ solution and then extracted with EtOAc (30 mL × 3). The organic layers were dried and concentrated. The residue was purified by column chromatography (DCM:MeOH = 20:1 to 15:1) to yield a colorless oil. Yield: 67%. $^1$H NMR (400 MHz, $CDCl_3$) δ 3.65 – 3.56 (m, 4H), 3.56 – 3.50 (m, 4H), 3.47 (t, $J$ = 6.7 Hz, 2H), 2.89 (t, $J$ = 4.7 Hz, 2H), 2.59 (s, 3H), 1.85 – 1.72 (m, 2H), 1.66 – 1.54 (m, 2H), 1.53 – 1.30 (m, 4H).

*Activation of glycol analogs* (**11a-c**). Glycol analogs **9a-c** (20 mmol) were added to a mixture of compound **10** (24 mmol) in 30 mL of DCM. Then, $Et_3N$ (30 mmol) was slowly added to the mixture at 0 °C. The mixture was then warmed to room temperature and stirred overnight. The reaction was diluted by adding 40 mL $H_2O$ and extracted with DCM (30 mL × 3). The organic layers were dried and concentrated. The residue was purified by column chromatography (PE:EtOAc = 4:1 to 3:1) to yield a yellowish oil. Yield: 81-92%. $^1$H NMR of **11c** (400 MHz, $CDCl_3$) δ 8.32 – 8.23 (m, 4H), 7.43 – 7.34 (m, 4H), 4.50 – 4.43 (m, 4H), 3.89 – 3.82 (m, 4H), 3.77 (s, 4H).

*Monosubstituted Halo ligands* (**12a-c**). Activated glycol analogs **11a-c** (10 mmol) and compound **8** (9 mmol) in 10 mL DMF were stirred at room temperature until reaction was complete, as monitored by TLC. The reaction was diluted by adding 20 mL of saturated aqueous $NH_4Cl$ and extracted with EtOAc (30 mL × 3). The organic layers were dried and concentrated. The residue was purified by column chromatography (PE: EtOAc = 1:1) to yield a yellowish oil. Yield: 55-69%. $^1$H NMR of **12c** (400 MHz, $CDCl_3$) δ 8.30 – 8.20 (m, 2H), 7.41 – 7.33 (m, 2H), 4.46 – 4.38 (m, 2H), 4.28 – 4.16 (m, 2H), 3.83 – 3.76 (m, 2H), 3.72 – 3.61 (m, 6H), 3.60 – 3.47 (m, 9H), 3.46 – 3.39 (m, 2H), 3.34 (t, $J$ = 5.3 Hz, 2H), 1.81 – 1.68 (m, 2H), 1.64 – 1.51 (m, 2H), 1.49 – 1.27 (m, 4H).

*Synthesis of dasatinib-Halo derivatives 2-5* (**DH2-5**). Dasatinib ligands were prepared as previously reported with some modification.[1] Briefly, dasatinib was activated by reacting with compound **10**, followed by the addition of diamine ligands **15a** and **15b** to form **16a** and **16b**. The reactions were diluted by adding 20 mL of saturated aqueous $NH_4Cl$ and extracted with EtOAc (30 mL × 3). Without further purification, they were reacted with monosubstituted halo ligands **12a-c** to afford **DH2-5** with diverse linker lengths **(**Off-white solid, yields: 20-35% over three steps).

*17-Chloro-4-oxo-3,8,11-trioxa-5-azaheptadecyl(2-(4-(6-((5-((2-chloro-6-methylphenyl)carbamoyl)thiazol-2-yl)amino)-2-methylpyrimidin-4-yl)piperazin-1-yl)ethyl) ethane-1,2-diyldicarbamate* (**DH2**). $^1$H NMR (400 MHz, $CD_3OD$) δ 8.19 (s, 1H), 7.36 (dd, $J$ = 7.3, 2.2 Hz, 1H), 7.30 – 7.19 (m, 2H), 6.01 (s, 1H), 4.24 (t, $J$ = 5.5 Hz, 2H), 4.16 (t, $J$ = 4.7 Hz, 4H), 3.78 – 3.63 (m, 8H), 3.63 – 3.51 (m, 8H), 3.47 (t, $J$ = 6.5 Hz, 2H), 3.29 (t, $J$ = 5.5 Hz, 2H), 3.23 (d, $J$ = 6.9 Hz, 4H), 2.72 (t, $J$ = 5.5 Hz, 2H), 2.65 (t, $J$ = 5.1 Hz, 4H), 2.49 (s, 3H), 2.34 (s, 3H), 1.81 – 1.70 (m, 2H), 1.64 – 1.53 (m, 2H), 1.51 – 1.37 (m, 4H). HRMS (ESI) calcd for $[C_{39}H_{57}Cl_2N_{10}O_9S]^+$ $[M+H]^+$ 911.3408, found 911.3416. HPLC: $t_R$ = 5.59 min, purity = 100%.

*17-Chloro-4-oxo-3,8,11-trioxa-5-azaheptadecyl(2-(4-(6-((5-((2-chloro-6-methylphenyl)carbamoyl)thiazol-2-yl)amino)-2-methylpyrimidin-4-yl)piperazin-1-yl)ethyl) ethane-1,2-diyldicarbamate* (**DH3**). $^1$H NMR (400 MHz, $CD_3OD$) δ 8.19 (s, 1H), 7.41 – 7.33 (m, 1H), 7.31 – 7.21 (m, 2H), 6.06 (s, 1H), 4.33 – 4.25 (m, 2H), 4.22 (m, 4H), 3.64 – 3.44 (m, 10H), 3.30 (t, $J$ = 5.5 Hz, 2H), 3.27 – 3.18 (m, 12H), 2.93 – 2.85 (m, 2H), 2.82 (m, 4H), 2.50 (s, 3H), 2.35 (s, 3H), 1.82 – 1.70 (m, 2H), 1.59 (m, 2H), 1.53 – 1.35 (m, 4H). HRMS (ESI) calcd for $[C_{41}H_{60}Cl_2N_{10}O_{10}SNa]^+$ $[M+Na]^+$ 977.3489, found 977.3497. HPLC: $t_R$ = 5.68 min, purity = 100%.

*23-Chloro-10-oxo-3,6,9,14,17-pentaoxa-11-azatricosyl(2-(4-(6-((5-((2-chloro-6-methylphenyl)carbamoyl)thiazol-2-yl)amino)-2-methylpyrimidin-4-yl)piperazin-1-yl)ethyl) ethane-1,2-diyldicarbamate* (**DH4**). $^1$H NMR (400 MHz, $CD_3OD$) δ 8.18 (s, 1H), 7.37 (dd, $J$ = 7.2, 2.2 Hz, 1H), 7.30 – 7.20 (m, 2H), 6.01 (s, 1H), 4.24 (t, $J$ = 5.5 Hz, 2H), 4.17 (t, $J$ = 4.7 Hz, 4H), 3.71 – 3.62 (m, 12H), 3.62 – 3.50 (m, 8H), 3.47 (t, $J$ = 6.6 Hz, 2H), 3.33 – 3.26 (m, 3H), 3.22 (s, 3H), 2.71 (t, $J$ = 5.6 Hz, 2H), 2.64 (t, $J$ = 4.9 Hz, 4H), 2.49 (s, 3H), 2.34 (s, 3H), 1.82 – 1.70 (m, 2H), 1.59 (m, 2H), 1.53 – 1.33 (m, 4H). HRMS (ESI) calcd for $[C_{43}H_{64}Cl_2N_{10}O_{11}SNa]^+$ $[M+Na]^+$ 1021.3751, found 1021.3751. HPLC: $t_R$ = 5.59 min, purity = 100%.

*23-Chloro-10-oxo-3,6,9,14,17-pentaoxa-11-azatricosyl(2-(4-(6-((5-((2-chloro-6-methylphenyl)carbamoyl)thiazol-2-yl)amino)-2-methylpyrimidin-4-yl)piperazin-1-yl)ethyl) pentane-1,5-diyldicarbamate* (**DH5**). $^1$H NMR (400 MHz, $CD_3OD$) δ 8.19 (s, 1H), 7.36 (dd, *J* = 7.3, 2.3 Hz, 1H), 7.30 – 7.19 (m, 2H), 6.00 (s, 1H), 4.22 (t, *J* = 5.5 Hz, 2H), 4.16 (dd, *J* = 6.1, 3.4 Hz, 4H), 3.73 – 3.61 (m, 12H), 3.61 – 3.50 (m, 8H), 3.47 (t, *J* = 6.6 Hz, 2H), 3.29 (t, *J* = 5.5 Hz, 2H), 3.11 (m, 4H), 2.69 (t, J = 5.5 Hz, 2H), 2.62 (t, *J* = 5.1 Hz, 4H), 2.48 (s, 3H), 2.34 (s, 3H), 1.82 – 1.70 (m, 2H), 1.64 – 1.26 (m, 12H). HRMS (ESI) calcd for $[C_{46}H_{70}Cl_2N_{10}O_{11}SNa]^+$ $[M+Na]^+$ 1063.4221, found 1063.4229. HPLC: $t_R$ = 5.60 min, purity = 100%.

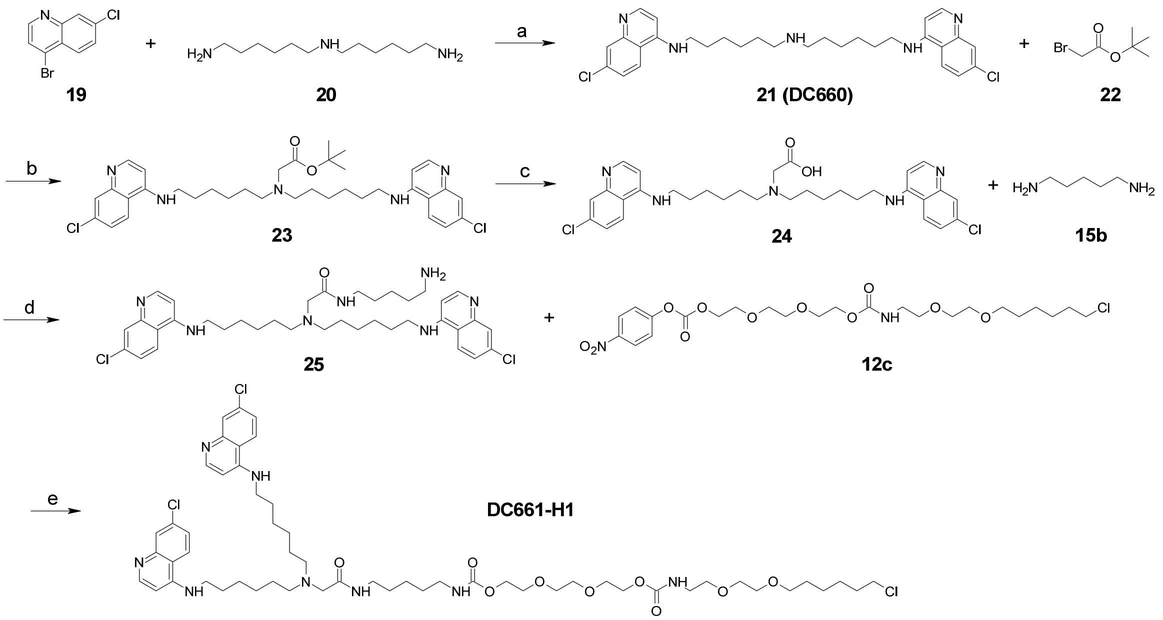


**Synthetic Scheme 3.** Synthesis of DC661-Halo 1 (**DC661-H1**). Reagents and conditions: (a) $Pd(OAc)_2$, BINAP, $K_3PO_4$, 1,4-Dioxane, 100 °C, 12 h; (b) $K_2CO_3$, DMF, 80 °C, 12 h; (c) TFA, DCM, RT, 1 h; (d) HATU, DIPEA, RT, 2 h; (e) DMF, RT, 2 h.

*$N^1$-(7-Chloroquinolin-4-yl)-$N^6$-(6-((7-chloroquinolin-4-yl)amino)hexyl)hexane-1,6-diamine* (**21, DC660**). A mixture of compound **19** (25 mmol), **20** (10 mmol), $Pd(OAc)_2$ (0.2 mmol), BINAP (0.4 mmol), and $K_3PO_4$ (30 mmol) in 20 mL of 1,4-dioxane was stirred at 100 °C overnight under an Ar atmosphere. The reaction was diluted by adding 20 mL of saturated aqueous $NH_4Cl$ and extracted with EtOAc (30 mL × 3). The organic layer was dried and concentrated. The residue was purified by column chromatography (DCM: MeOH = 10:1 to 5:1) to yield a white solid. Yield: 83%. $^1$H NMR (400 MHz, $CD_3OD$) δ 8.36 (d, *J* = 5.6 Hz, 2H), 8.12 (d, *J* = 9.0 Hz, 2H), 7.79 (d, *J* = 2.2 Hz, 2H), 7.41 (dd, *J* = 9.0, 2.2 Hz, 2H), 6.53 (d, *J* = 5.7 Hz, 2H), 3.39 (t, *J* = 7.2 Hz, 4H), 2.76 – 2.65 (m, 4H), 1.84 – 1.72 (m, 4H), 1.65 – 1.40 (m, 12H).

*Tert-butyl bis(6-((7-chloroquinolin-4-yl)amino)hexyl)glycinate* (**23**). A mixture of compound **21** (10 mmol), **22** (12 mmol), and $K_2CO_3$ (15 mmol) in 20 mL of DMF was stirred at 80 °C overnight under an Ar atmosphere. The reaction was diluted by adding 20 mL of saturated aqueous $NH_4Cl$ and extracted with EtOAc (30 mL × 3). The organic layer was dried and concentrated. The residue was purified by column chromatography (DCM: MeOH = 50:1 to 20:1) to yield an off-white solid. Yield: 79%. $^1$H NMR (400 MHz, $CDCl_3$) δ 8.40 (d, *J* = 5.9 Hz, 2H), 8.19 (d, *J* = 9.0 Hz, 2H), 8.00 (d, *J* = 2.1 Hz, 2H), 7.35 (dd, *J* = 9.0, 2.1 Hz, 2H), 6.83 (s,

2H), 6.44 (d, $J$ = 5.9 Hz, 2H), 3.40 (q, $J$ = 6.7 Hz, 4H), 3.20 (s, 2H), 2.54 (t, $J$ = 7.0 Hz, 4H), 1.76 (m, 4H), 1.52 – 1.24 (m, 21H).

*Synthesis of DC661-Halo derivative 1* (**DC661-H1**). The deprotection of compound **23** (5 mmol) was carried out by adding 2 mL of TFA. The TFA was then evaporated under vacuum to afford compound **24**. The residue was used directly for the next step. A mixture of **24**, **15b** (5 mmol), HATU (6 mmol), and DIPEA (20 mmol) in 10 mL of DMF was stirred at room temperature for 1 h. The reaction was then diluted by adding 20 mL of saturated aqueous $NH_4Cl$ and extracted with EtOAc (30 mL × 3). The organic layer was dried and concentrated. The residue was further reacted with **12c** (6 mmol) for 2 h. The reaction was again diluted by adding 20 mL of saturated aqueous $NH_4Cl$ and extracted with EtOAc (30 mL × 3). The organic layer was dried and concentrated. The residue was purified by column chromatography (DCM: MeOH = 30:1 to 20:1) to yield a yellowish solid. Yield: 43% over 3 steps. $^1H$ NMR (400 MHz, $CD_3OD$) δ 8.37 (d, $J$ = 6.3 Hz, 2H), 8.24 (d, $J$ = 9.0 Hz, 2H), 7.80 (d, $J$ = 2.1 Hz, 2H), 7.52 (dd, $J$ = 9.1, 2.1 Hz, 2H), 6.67 (d, $J$ = 6.3 Hz, 2H), 4.16 (dq, $J$ = 12.8, 4.7 Hz, 6H), 3.73 – 3.42 (m, 20H), 3.29 (q, $J$ = 5.8 Hz, 4H), 3.21 (t, $J$ = 7.3 Hz, 2H), 3.12 (s, 2H), 3.10 – 2.99 (m, 2H), 2.56 (t, $J$ = 7.4 Hz, 4H), 1.67 – 1.23 (m, 28H). $^{13}C$ NMR (151 MHz, $CD_3OD$) δ 172.34, 157.45, 157.45, 153.48, 153.48, 147.00, 147.00, 143.81, 143.81, 136.98, 136.98, 125.75, 125.75, 123.68, 123.68, 122.84, 122.84, 116.47, 116.47, 98.24, 98.24, 72.29, 70.79, 70.78, 70.18, 70.15, 70.00, 69.84, 69.75, 69.52, 69.18, 69.16, 63.67, 63.56, 60.81, 57.59, 55.15, 44.31, 44.29, 42.97, 40.28, 40.21, 38.46, 32.32, 29.12, 29.10, 28.83, 27.80, 26.75, 26.63, 26.58, 26.31, 25.05, 23.73, 21.12. HRMS (ESI) calcd for $[C_{55}H_{83}Cl_3N_8O_9Na]^+$ $[M+Na]^+$ 1127.5246, found 1127.5236.

**A**

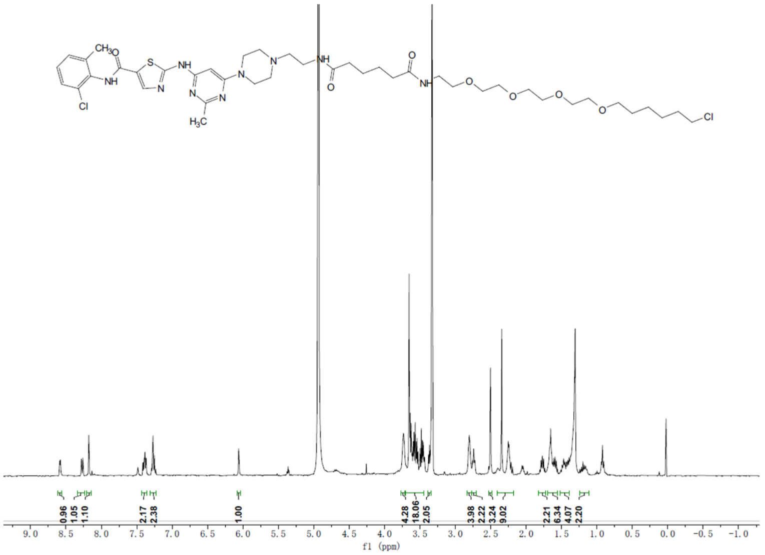


**B**

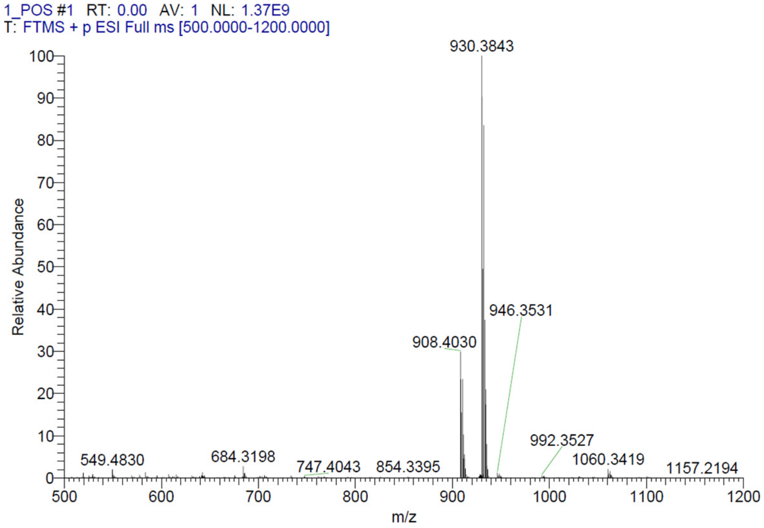


**Representative spectra of DH1.** (A) $^1$H NMR (B) HRMS

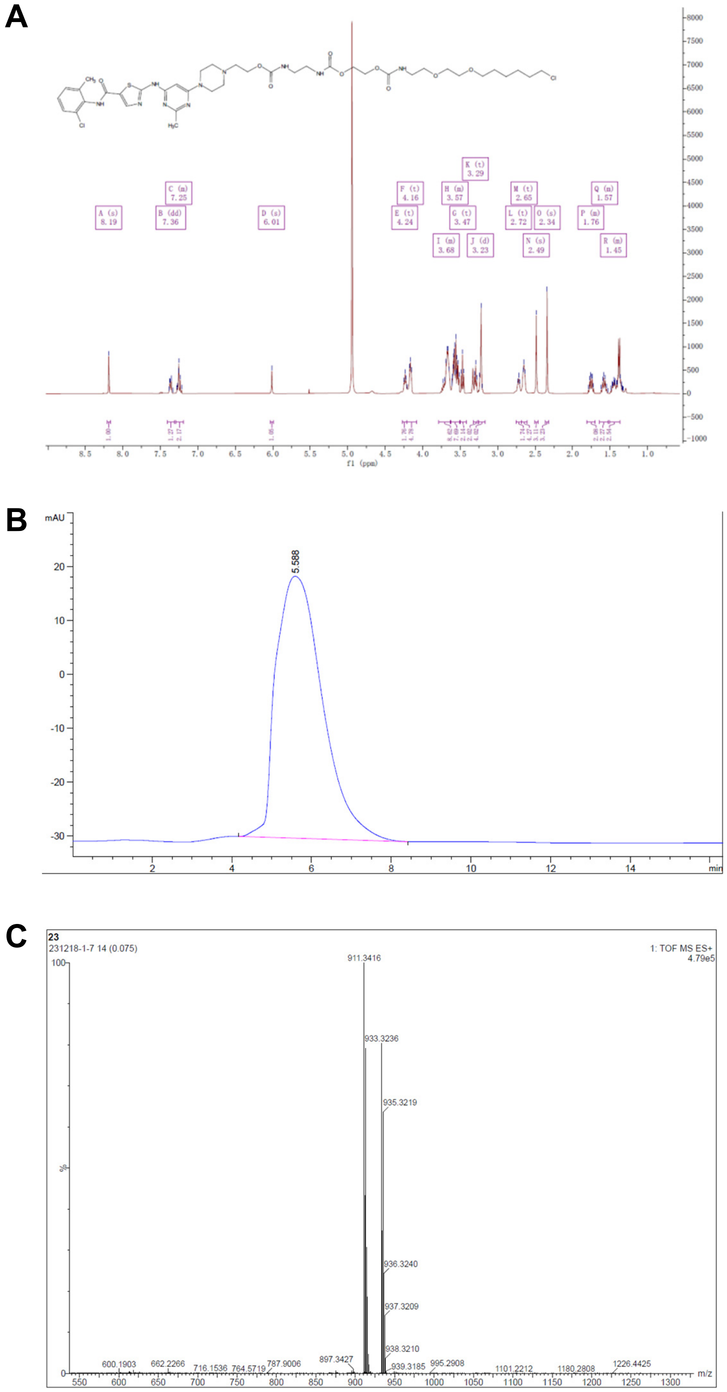


**Representative spectra of DH2.** (A) $^{1}$H NMR (B) HPLC (C) HRMS

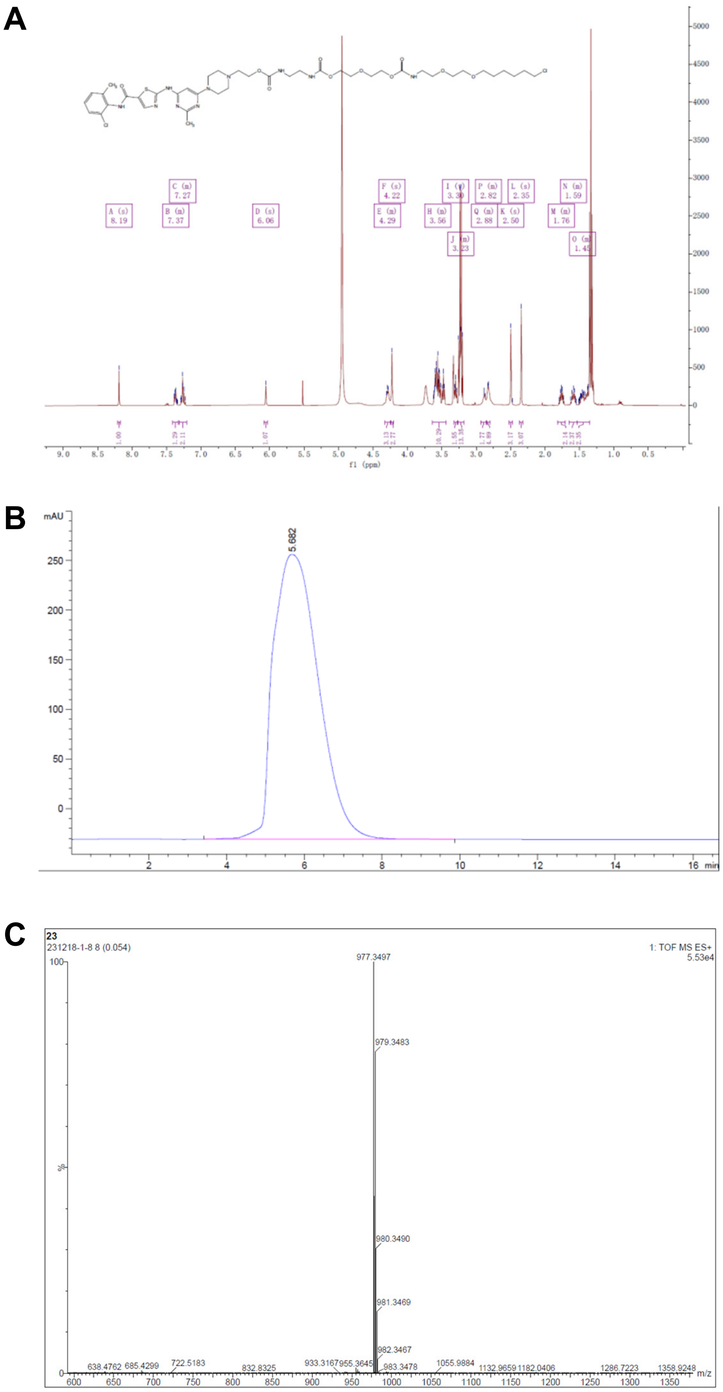


**Representative spectra of DH3.** (A) $^{1}$H NMR (B) HPLC (C) HRMS

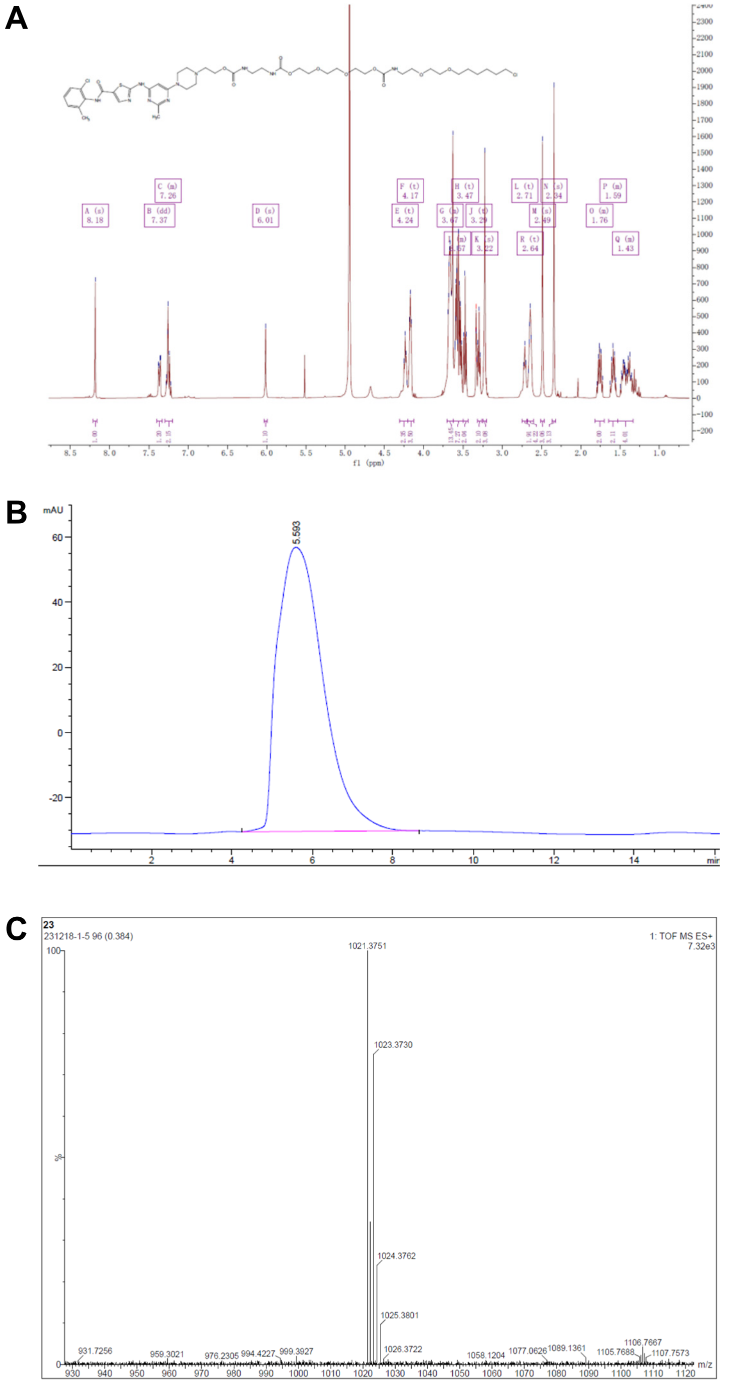

**Representative spectra of DH4.** (A) $^1$H NMR (B) HPLC (C) HRMS

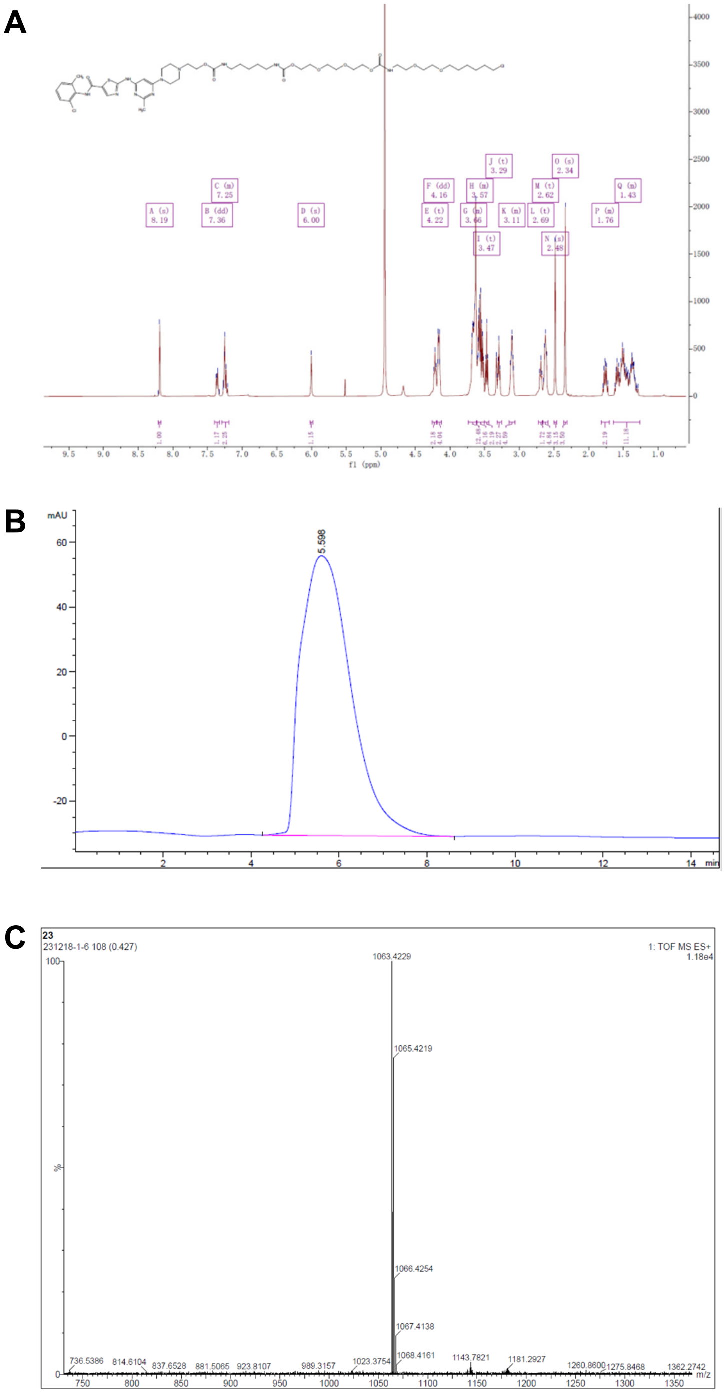


**Representative spectra of DH5.** (A) $^{1}$H NMR (B) HPLC (C) HRMS

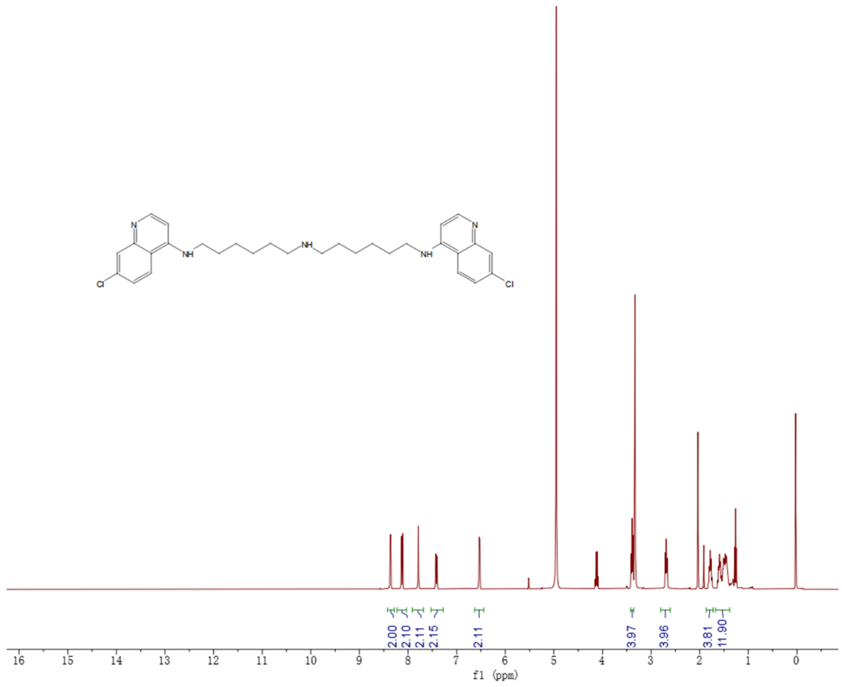


**Representative spectra.** $^{1}$H NMR of DC660

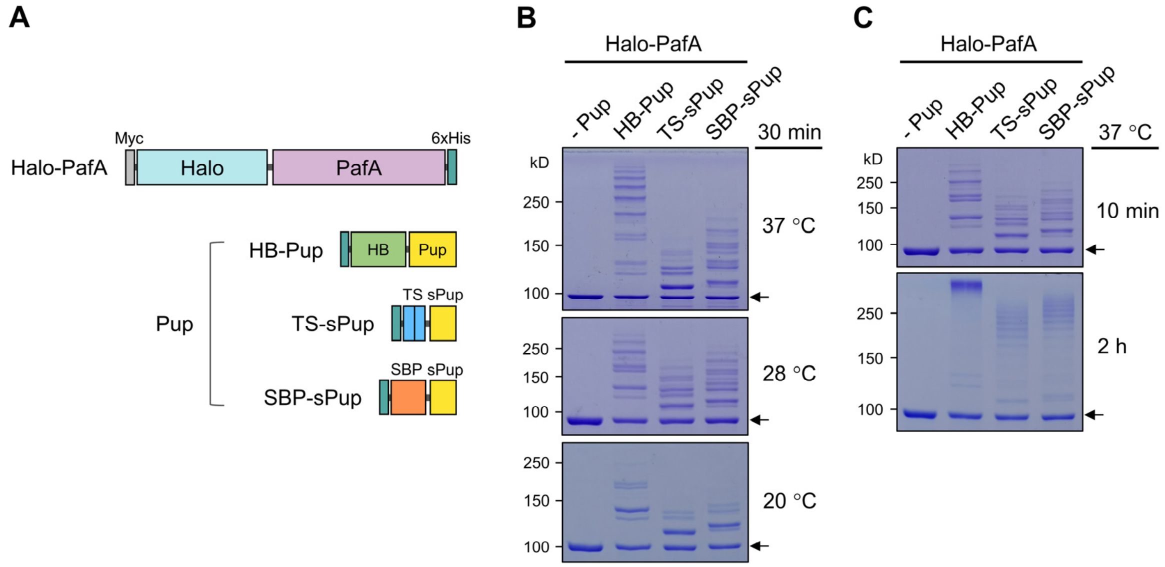


**Figure 1—figure supplement 1.** PafA fused with HaloTag exhibits high activity across a wide temperature range. (**A**) Schematic representation of Halo-PafA and three Pup substrates (HB-Pup, TS-sPup, SBP-sPup). HB, TS, and SBP refer to 6×His and BCCP, twin-STII (Strep-tag II), and streptavidin binding peptide, respectively. (**B**) In vitro self-pupylation of Halo-PafA (1 μM) with HB-Pup, TS-sPup, or SBP-sPup (10 μM) at 37 °C, 28 °C, and 20 °C for 30 min. (**C**) Polypupylated Halo-PafAs were observed as high molecular weight bands after 2 h pupylation reaction with three Pup substrates (HB-Pup, TS-sPup, SBP-sPup) at 37 °C. Arrows indicate the Halo-PafA band without pupylation.

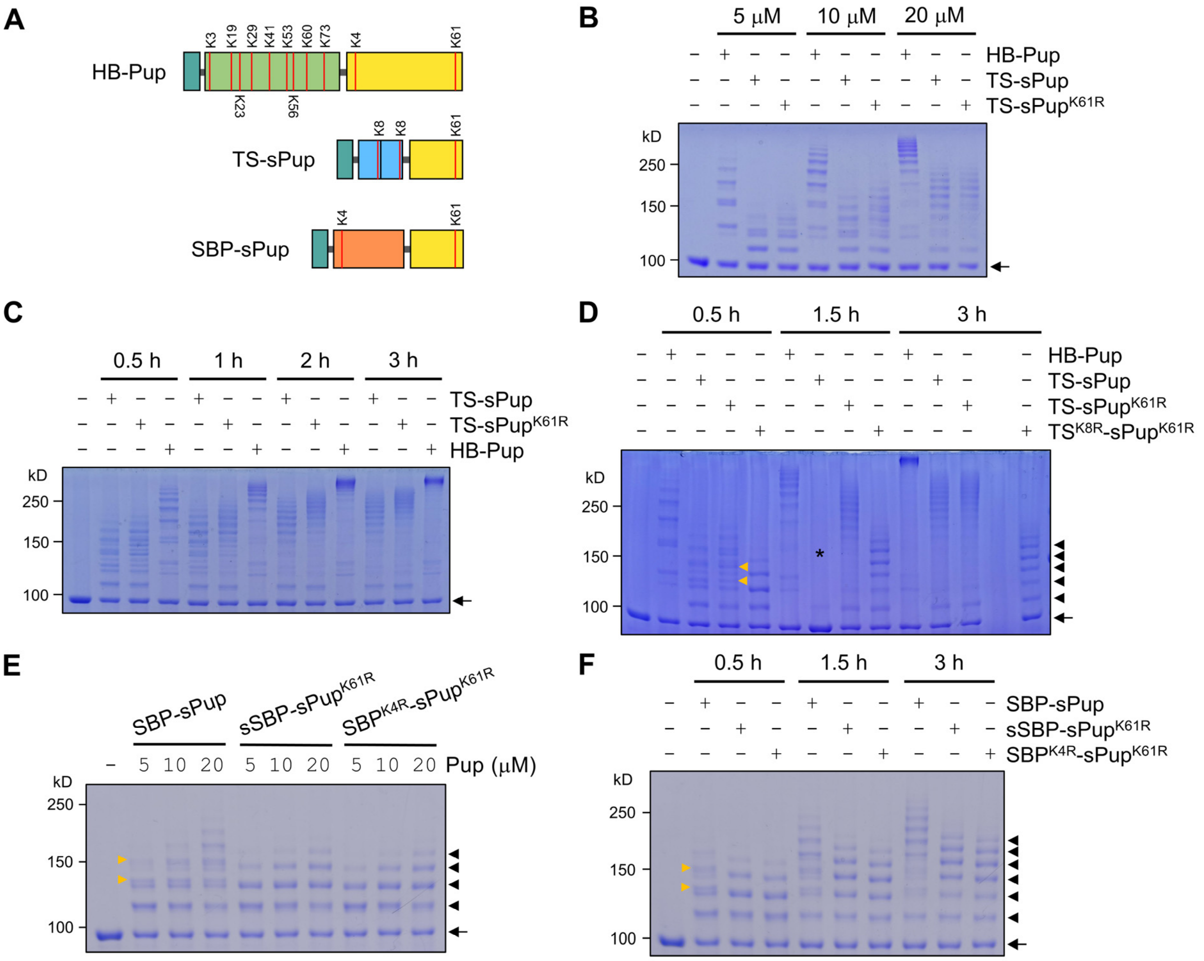


**Figure 1—figure supplement 2.** Lysine-mutated Pup substrates bypass polypupylation while retaining activities comparable to wild-type Pup. (**A**) A schematic shows the lysine residues in HB-Pup, TS-sPup, and SBP-sPup. (**B**) In a pupylation assay, Halo-PafA exhibits dose-dependent self-pupylation with different concentrations of HB-Pup, TS-sPup, or TS-sPup$^{K61R}$ for 30 min. (**C**) Time-dependent polypupylation becomes evident, particularly after 1 h incubation, as seen by high molecular weight bands. (**D**) Removing all lysine residues from TS-sPup completely prevents polypupylation. The asterisk indicates a possible experimental error causing a failed pupylation. (**E**) Eliminating lysine residues in SBP and sPup completely abolishes polypupylation. Halo-PafA was incubated with various concentrations of SBP-sPup, sSBP-sPup$^{K61R}$, or SBP$^{K4R}$-sPup$^{K61R}$ for 30 min. (**F**) Longer incubations do not induce polypupylation but enhance multipupylation in the cases of sSBP-sPup$^{K61R}$ and SBP$^{K4R}$-sPup$^{K61R}$. All reactions were performed with 1 µM of Halo-PafA and 10 µM of HB-Pup, TS-sPup, or TS-sPup$^{K61R}$ at 37 °C. Yellow arrowheads point to polypupylated bands which are not detected in polypupylation-free sPup substrates. Black arrowheads and arrows indicate multipupylated bands and Halo-PafA, respectively.

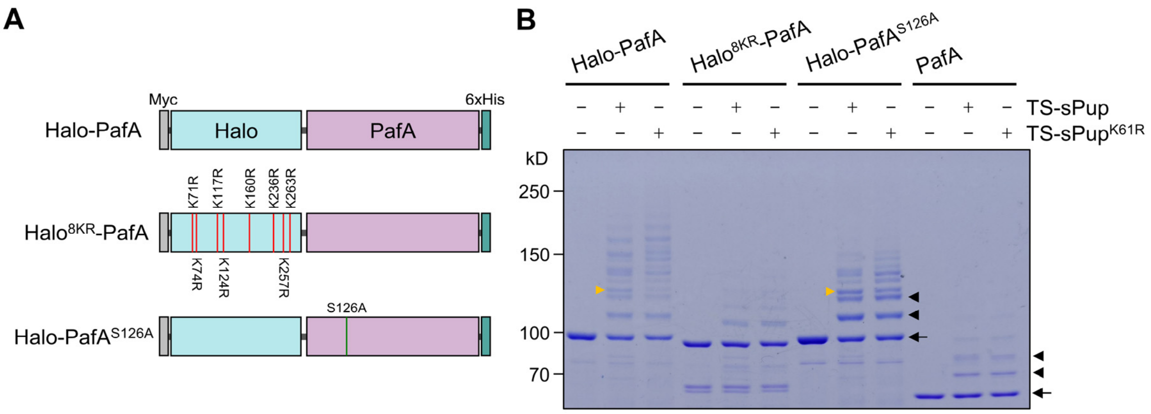


**Figure 1—figure supplement 3.** Optimization of Halo-PafA. (**A**) Schematic illustration of Halo-PafA derivatives: $Halo^{8KR}$-PafA, which includes eight lysine-to-arginine mutations to abolish self-pupylation, and Halo-$PafA^{S126A}$, which contains a serine-to-alanine mutation at position 126 to decrease depupylase activity of PafA. (**B**) In vitro self-pupylation results show that $Halo^{8KR}$-PafA exhibits minimal pupylation levels, similar to PafA alone. Halo-$PafA^{S126A}$ demonstrates reduced self- and polypupylation while enhancing multipupylation. Reactions were conducted with 1 μM of a Halo-PafA derivative and 10 μM of TS-sPup or TS-$sPup^{K61R}$ at 37 °C for 30 min. Yellow arrowheads indicate polypupylated bands, while black arrowheads and arrows indicate multipupylated bands and Halo-PafA, respectively.

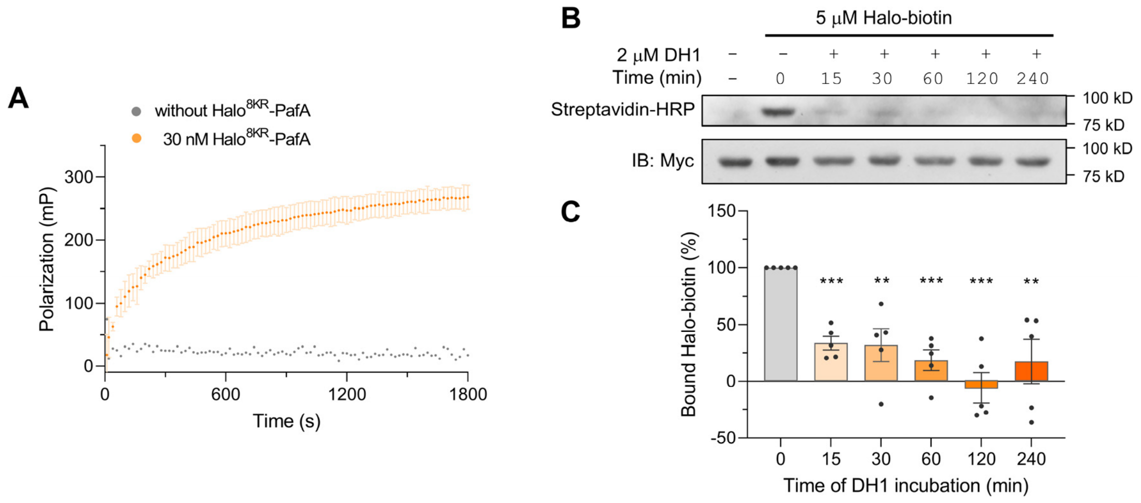


**Figure 2—figure supplement 1.** Halo$^{8KR}$-PafA demonstrates efficient labeling with HaloTag ligands. (**A**) Fluorescence polarization (FP) assay using 2 nM of HaloTag Alexa Fluor-488 (Halo-AF488) at 25 °C. Data are shown as mean ± s.d. (**B**)Western blot analysis for the Halo-Biotin competition assay. Halo-Biotin (5 μM) was incubated with 1 μM of Myc-Halo$^{8KR}$-PafA at 37 °C for 1 h after a preincubation with 2 μM of DH1 for the indicated time period. Levels of bound Halo-Biotin were assessed by streptavidin-HRP. (**C**) Quantitative analysis of western blot results shown in (**B**). $n = 5$. Data are shown as mean ± s.e.m. $P$ values were calculated by an unpaired two-sided $t$-test. $^{**}P < 0.01$, $^{***}P < 0.001$; versus no DH1 competition.

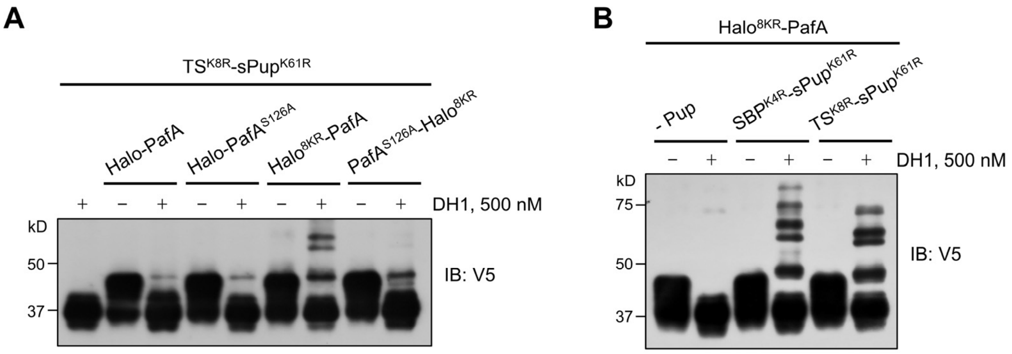


**Figure 2—figure supplement 2.** Optimized Halo-PafA and Pup substrates efficiently promote pupylation in vitro in a proximity-dependent manner. (**A**) Comparison of proximity-tagging by different Halo-PafA derivatives on in vitro pupylation using $TS^{K8R}$-$sPup^{K61R}$. (**B**) Comparison of different Pup substrates, $SBP^{K4R}$-$sPup^{K61R}$ and $TS^{K8R}$-$sPup^{K61R}$, on in vitro pupylation. All reactions were conducted with 1 μM of a Halo-PafA derivative, 10 μM of a sPup substrate, and 0.5 μM of a purified short SRC with a 2×V5 tag, SRC(247-536)-2×V5, at 37 °C for 30 min. Pupylation levels of SRC were assessed by immunoblot (IB) using an anti-V5 antibody.

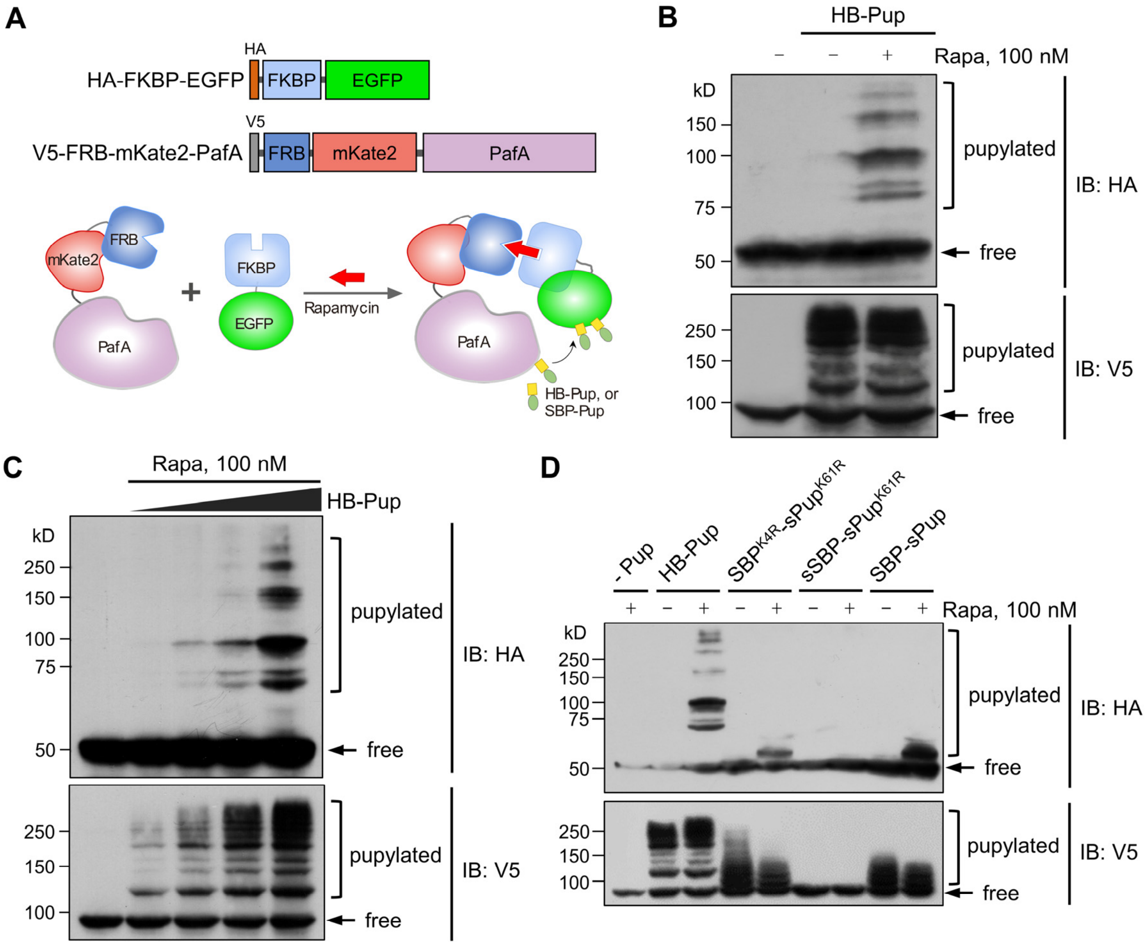


**Figure 3—figure supplement 1.** Test of small molecule-induced proximity-tagging by PafA in HEK293T cells. (**A**) Schematic illustration of plasmid constructs and a model of proximity-tagging by PafA. In the presence of rapamycin, HA-FKBP-EGFP and V5-FRB-mKate2-PafA heterodimerize via rapamycin, facilitating proximity for PafA to pupylate HA-FKBP-EGFP. (**B**) V5-FKBP-EGFP is highly pupylated only in the presence of rapamycin, while FRB-mKate2-PafA undergoes substantial self-pupylation regardless of rapamycin treatment. (**C**) The pupylation levels of the target, HA-FKBP-EGFP, and the self-pupylation of V5-FRB-mKate2-PafA increase in an HB-Pup dose-dependent manner. (**D**) Evaluation of different Pup substrates on proximity-tagging and self-pupylation. (**B**-**D**) HEK293T cells were co-transfected with HA-FKBP-EGFP, V5-FRB-mKate2-PafA, and an indicated Pup substrate. Twenty-four hours post-transfection, the cells were treated with 100 nM rapamycin for an additional 24 h. Pupylation levels of HA-FKBP-EGFP and V5-FRB-mKate2-PafA were assessed by immunoblot (IB) using antibodies against HA-tag and V5-tag, respectively.

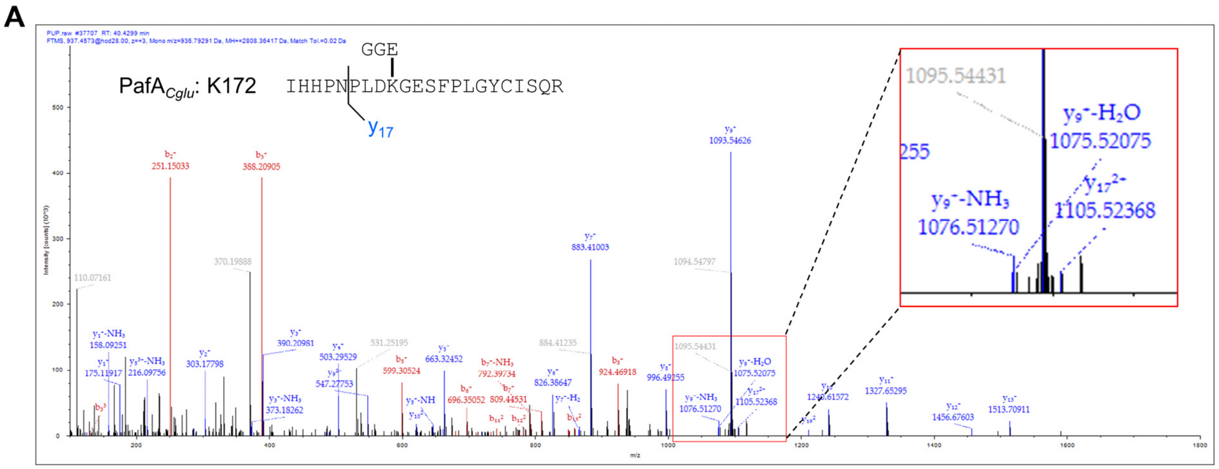


1H + 1909.948 ($y_{17}^{+}$) + 57.021 (carbamidomethylation on C18) + 243.086 (GGE))/2=1105.5275= $y_{17}^{2+}$ (1105.52368)

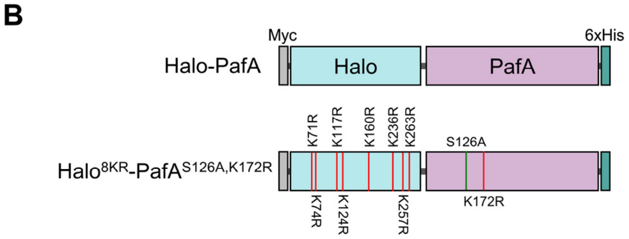


**Figure 3—figure supplement 2.** Identification of self-pupylated lysine residues in PafA. (**A**) Mass spectrometry analysis identified K172 as the most frequently pupylated lysine in PafA. An inset provides an enlarged view of the Nano-LC-MS/MS mass spectrum. (**B**) Constructs of Halo-PafA and Halo$^{8KR}$-PafA$^{S126A,K172R}$ containing 8KR mutations in HaloTag and mutations of S126A and K172R in PafA.

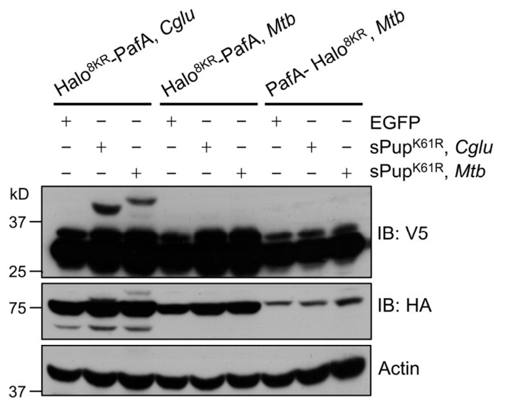


**Figure 3—figure supplement 3.** PafA from *Cglu* outperforms PafA from *Mtb* in the POST-IT system in live cells. HEK293T cells were co-transfected with one of the HA-Halo-PafA derivatives and a sPup substrate as shown above, along with SRC(247-536)-2×V5 as a target. Twenty-four hours later, cells were incubated with 250 nM DH1 for an additional 24 h. Immunoblot (IB) analysis reveals that PafA from *Cglu* exhibits robust labeling with $sPup^{K61R}$, whether from *Cglu* or *Mtb*, whereas PafA from *Mtb* shows no detectable labeling.

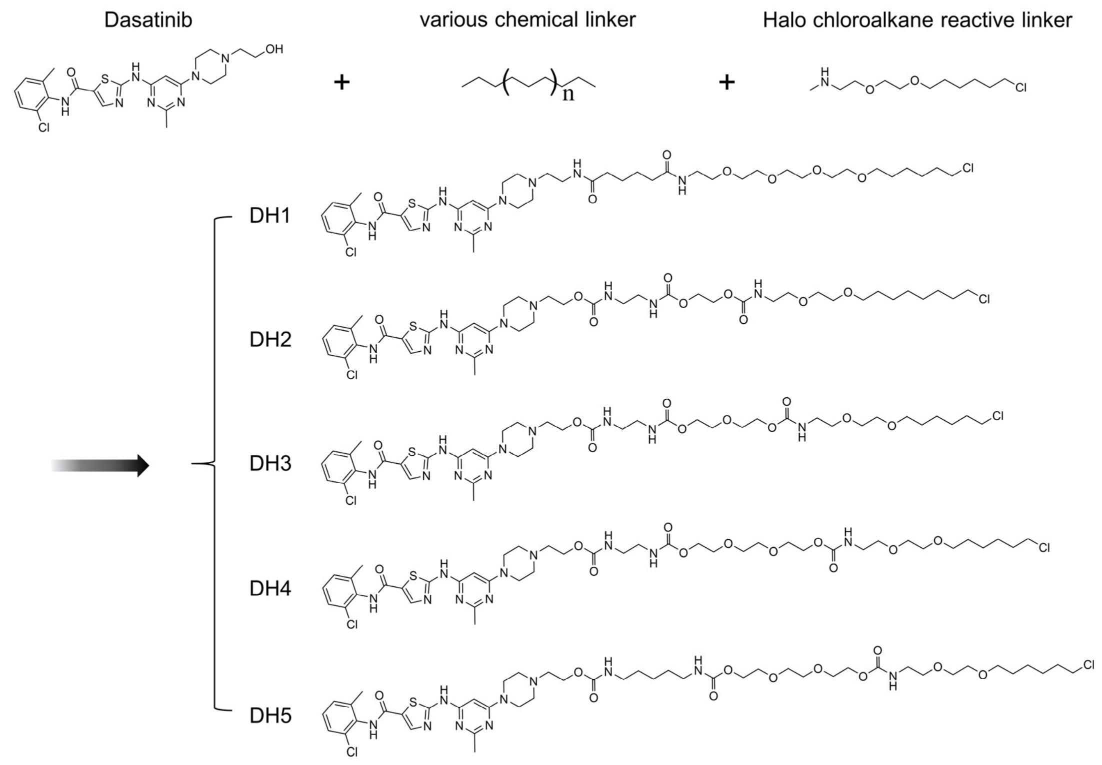


**Figure 4—figure supplement 1.** Chemical structures of dasatinib and its HTL derivatives with varied linkers.

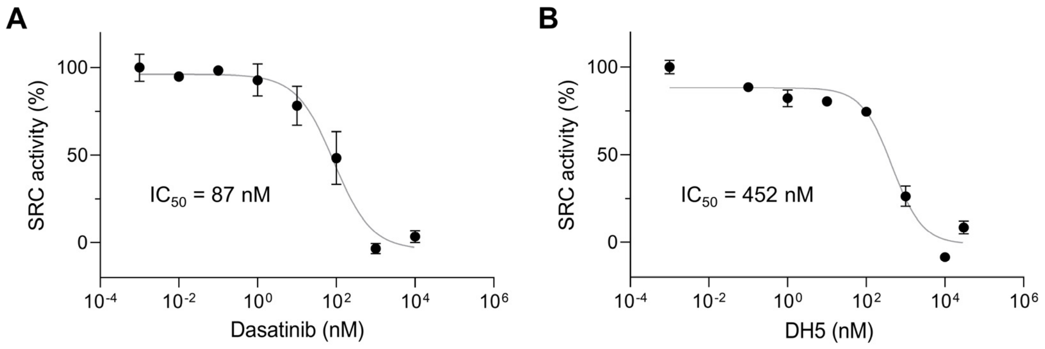


**Figure 4—figure supplement 2.** DH5 maintained robust kinase inhibitory activity. (**A, B**) SRC kinase activity was measured in vitro to assess inhibitory activities of dasatinib (**A**) and DH5 (**B**). A purified, catalytically active SRC(251-536), at a concentration of 0.8 μM, was incubated with a serial dilution of either dasatinib or DH5. The $IC_{50}$ for each compound was determined from their dose-response curves. Data are shown as mean ± s.e.m.

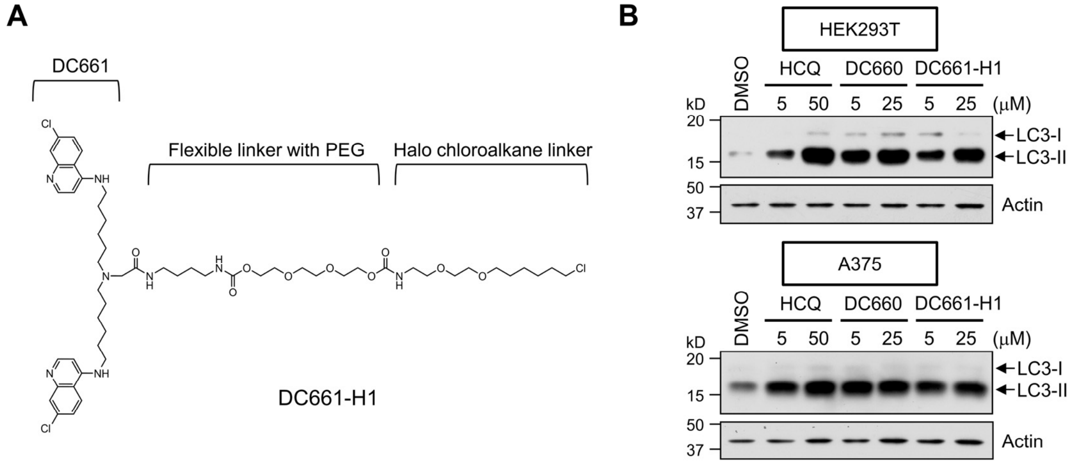


**Figure 6—figure supplement 1.** A DC661-HTL derivative, DC661-H1, is highly active as an autophagy inhibitor. (**A**) The chemical structure of DC661-H1, annotated with key functional groups. (**B**) Autophagy inhibition by DC661-H1 was assessed through the increased levels of the LC3-II band by western blot using an anti-LC3B antibody. DC661-H1 significantly increased the levels of LC3-II in both HEK293T and A375 cells, comparably to HCQ and DC660, an analog of DC661 that lacks a methyl group at the central linker nitrogen. Cells were treated with the indicated concentration of each compound for 12 h prior to western blot analysis.

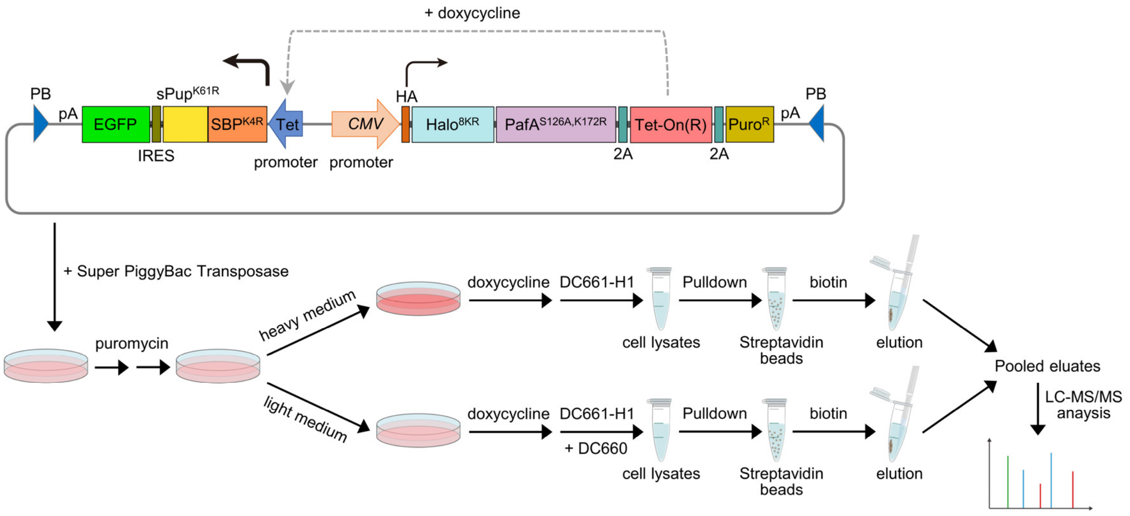


**Figure 6—figure supplement 2.** Schematic of the experimental flow for applying POST-IT in SILAC. A PiggyBac (PB) plasmid construct containing the POST-IT system was co-transfected with the Super PiggyBac Transposase expression vector into HEK293T cells. Four days later, the cells were treated with 5 µg/ml puromycin for approximately two weeks. Media containing puromycin were changed every 3–4 days. Cells were divided into two groups and treated with either heavy (K8R10) or light medium (K0R0). The expression of POST-IT was induced by the addition of 100 ng/ml doxycycline. Two days later, 500 nM DC661-H1 was added to the heavy medium, or 500 nM DC661-H1 and 5 µM DC660 to the light medium. After 24 h of incubation, cell lysates were prepared, pulled down by streptavidin magnetic beads, washed sequentially with wash buffer A and then wash buffer B, and eluted with 5 mM biotin and 10 µM DC660. The eluates were combined and precipitated using TCA/DOC, separated by Tricine-SDS-PAGE, and analyzed by LC-MS/MS. Refer to the methods section for more details.

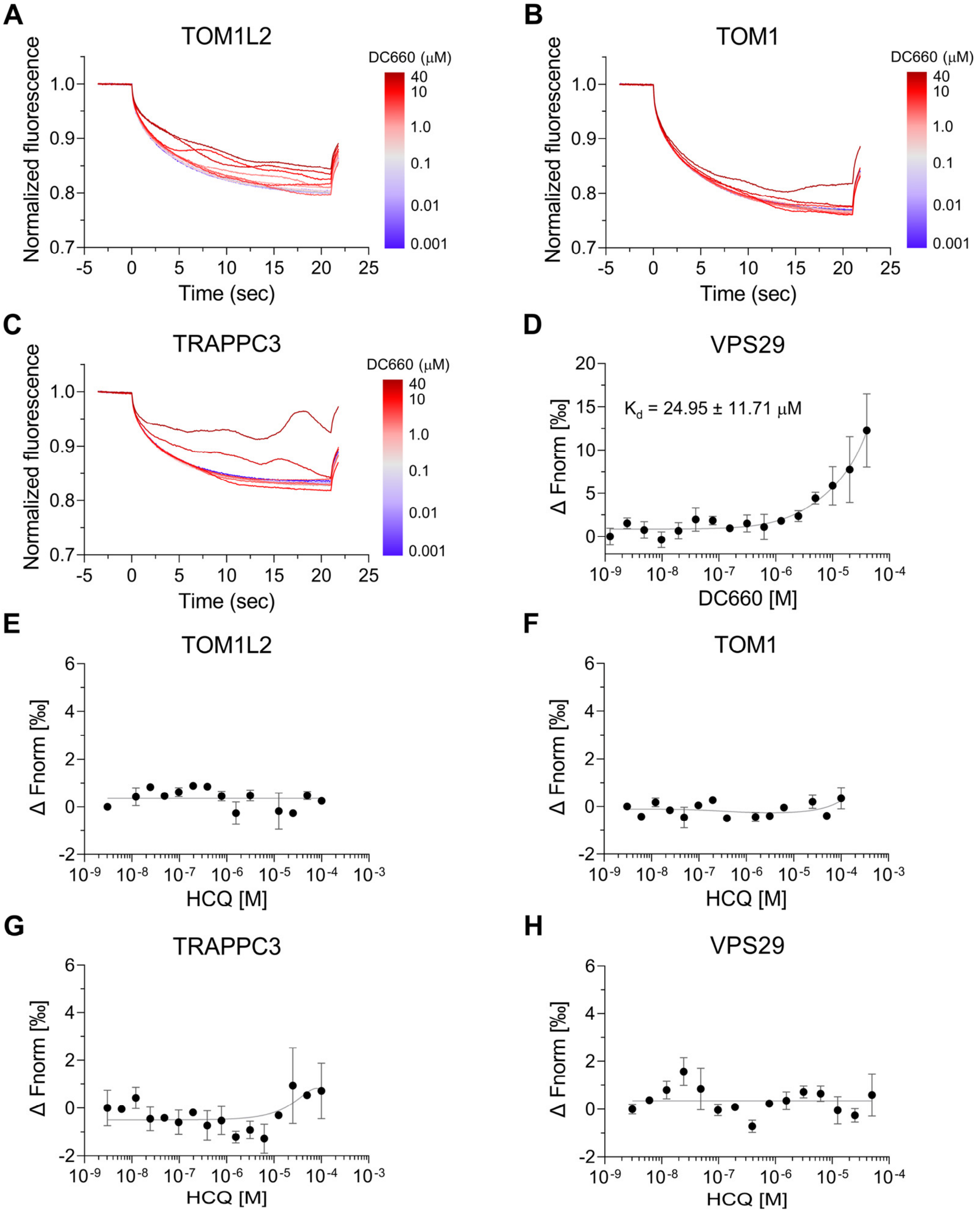


**Figure 6—figure supplement 3.** MST binding analysis of candidate proteins identified with DC661-H1. (**A-C**) Thermophoretic movement of DC660 bound to purified target proteins labeled with Cy5 dye. TOM1L2, TOM1, and TRAPPC3 exhibited ligand-induced aggregation, suggesting their interaction with DC660. (**D-H**) Dose-response curves of normalized fluorescence changes for target proteins binding. VPS29 demonstrated substantial affinity to DC660 with a $K_d$ of 24.95 ± 11.71 μM (**D**), but showed no binding to HCQ (**H**). TOM1L2, TOM1, and TRAPPC3 did not show detectable binding to HCQ (**E-H**). Data are shown as mean ± s.d.

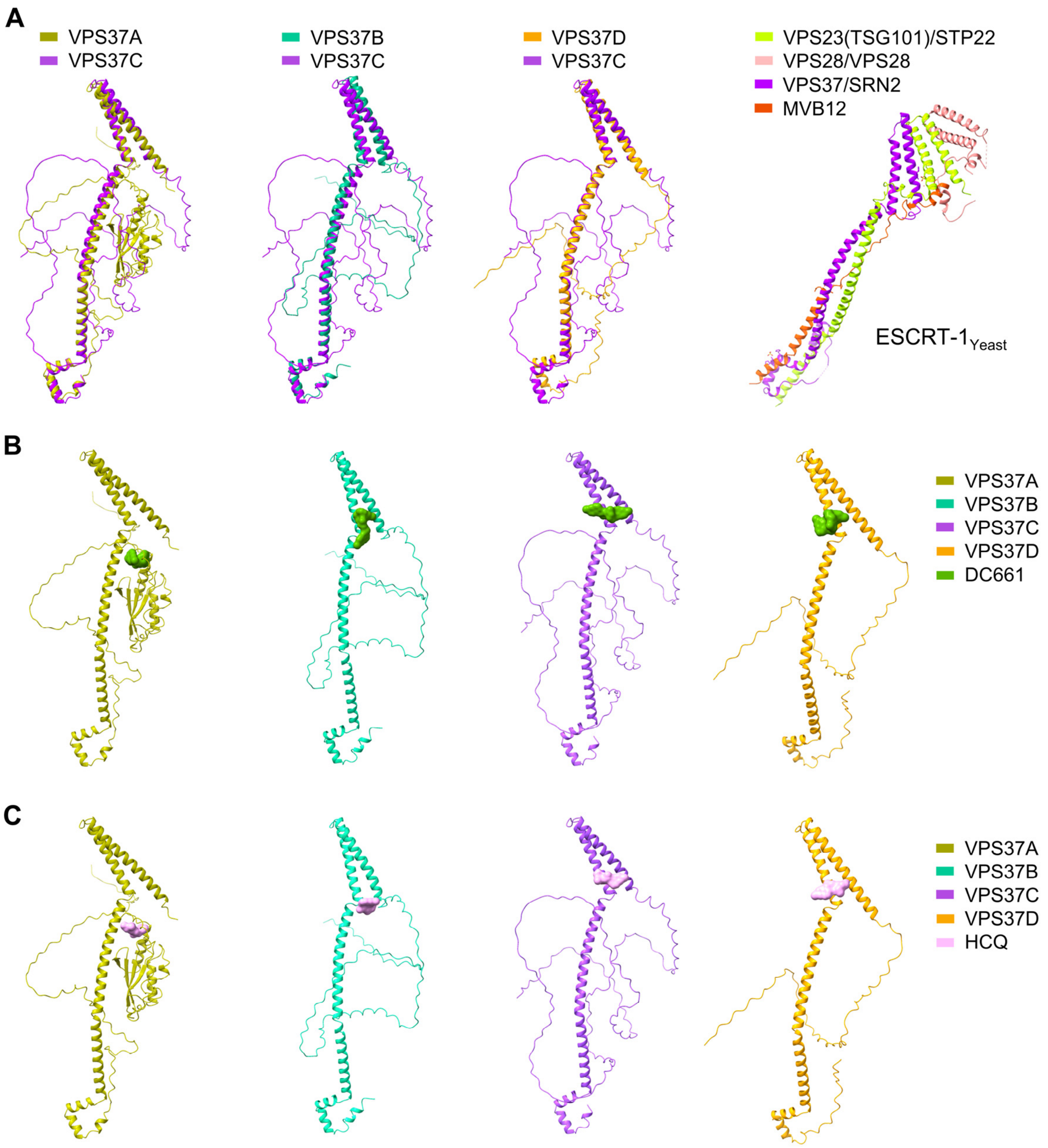


**Figure 6—figure supplement 4.** Molecular docking binding poses of DC661 and HCQ with the VPS37 family. (**A**) Structural superposition of VPS37C with other members of the VPS37A/B/D family reveals high structural similarities within the VPS37 family. The crystal structure of the ESCRT-1 complex from yeast (2P22) is shown on the far right. STP22 and SRN2 are yeast orthologs of VPS23(TSG101) and VPS37, respectively. (**B**) Molecular docking binding poses of DC661 with the VPS37 family. (**C**) Molecular docking binding poses of HCQ with the VPS37 family.

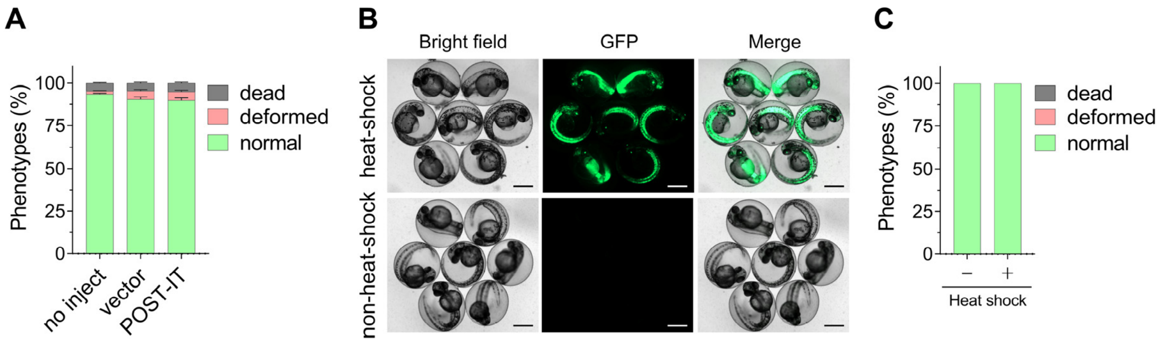


**Figure 7—figure supplement 1.** Expression of POST-IT does not cause toxicity in zebrafish. (**A**) Quantitative analysis shows that embryos injected with POST-IT plasmid exhibit no signs of toxicity. Embryos were injected at 1-cell stage, and phenotypes were assessed 24 h later. $n = 3$. Data are shown as mean ± s.e.m. (**B**) Representative images of transgenic zebrafish embryos expressing heat-shock-inducible POST-IT show no obvious toxicity after heat shock. EGFP images confirm robust expression of POST-IT. Healthy embryos at 1 dpf were treated at 39 °C for 30 min and imaged at 2 dpf. Scale bar, 500 µm. (**C**) Quantitative data related to (B) indicate that POST-IT expression following heat treatment does not cause toxicity. $n = 3$. Data are shown as mean ± s.e.m.

**Data S1. (separate file)**
Proteomics result for DH5

**Data S2. (separate file)**
SILAC result for DC661-H1
